# Genome-wide co-localization of RNA-DNA interactions and fusion RNA pairs

**DOI:** 10.1101/472019

**Authors:** Zhangming Yan, Norman Huang, Weixin Wu, Weizhong Chen, Yiqun Jiang, Jingyao Chen, Xuerui Huang, Xingzhao Wen, Jie Xu, Qiushi Jin, Kang Zhang, Zhen Chen, Shu Chien, Sheng Zhong

## Abstract

Fusion transcripts are used as biomarkers in companion diagnoses. Although more than 15,000 fusion RNAs have been identified from diverse cancer types, few common features have been reported. Here, we compared 16,410 fusion transcripts detected in cancer (from a published cohort of 9,966 tumor samples of 33 cancer types) with genome-wide RNA-DNA interactions mapped in two normal, non-cancerous cell types (using iMARGI, an enhanced version of the MARGI [<u>Ma</u>pping <u>R</u>NA-<u>G</u>enome <u>I</u>nteractions assay]). Among the top 10 most significant RNA-DNA interactions in normal cells, 5 co-localized with the gene pairs that formed fusion RNAs in cancer. Furthermore, throughout the genome, the frequency of a gene pair to exhibit RNA-DNA interactions is positively correlated with the probability of this gene pair to present documented fusion transcripts in cancer. To test whether RNA-DNA interactions in normal cells are predictive of fusion RNAs, we analyzed these in a validation cohort of 96 lung cancer samples using RNA-seq. 37 out of 42 fusion transcripts in the validation cohort were found to exhibit RNA-DNA interactions in normal cells. Finally, by combining RNA-seq, single-molecule RNA FISH, and DNA FISH, we detected a cancer sample with EML4-ALK fusion RNA without forming the EML4-ALK fusion gene. Collectively, these data suggest a novel RNA-poise model, where spatial proximity of RNA and DNA could poise for the creation of fusion transcripts.

---

Fusion transcripts are associated with diverse cancer types and have been proposed as diagnostic biomarkers (1–3). Companion tests and targeted therapies have been developed to identify and treat fusion-gene defined cancer subtypes (2, 4). Efforts of detection of fusion transcripts have primarily relied on analyses of RNA sequencing (RNA-seq) data (1, 3, 5–8). A recent study analyzed 9,966 RNA-seq datasets across 33 cancer types from The Cancer Genome Atlas (TCGA) and identified more than 15,000 fusion transcripts (4).

Despite the large number of gene pairs in identified fusion transcripts, it remains formidable to predict what unreported pair of genes may form a new fusion transcript. Recent analyses could not identify any distinct feature of fusion RNA forming gene pairs (9). Here, we report a characteristic pattern of the genomic locations of the gene pairs involved in fusion transcripts.

Chromatin-associated RNAs (caRNAs) provide an additional layer of epigenomic information in parallel to DNA and histone modifications (10). The recently developed MARGI (Mapping RNA-Genome Interactions) technology enabled identification of diverse caRNAs and the respective genomic interacting locations of each caRNA (6). In this work, we used an improved MARGI experimental pipeline to map RNA-DNA interactions in human embryonic kidney (HEK) and human foreskin fibroblast (HFF) cells. The detected RNA-DNA interactions often appeared on the gene pairs involved in cancer-derived fusion transcripts. The wide-spread RNA-DNA interactions on the gene pairs involved in fusion transcripts suggest a model, wherein the RNA of gene 1 by interacting with the genomic sequence of gene 2 is poised for being spliced into gene 2’s nascent transcript and thus creating a fusion transcript. Consistent with this model, we identified an RNA fusion in a new cancer sample that does not involve the creation of a fusion gene.

## Results

### Characteristics of genome-wide RNA-DNA interaction maps

To systematically characterize caRNAs and their genomic interaction locations, we developed iMARGI, an enhanced version of the MARGI assay (10). The main difference between iMARGI and MARGI is that iMARGI carries out the ligation steps *in situ* (Fig. 1A), whereas MARGI performs these ligation steps on streptavidin beads. We applied iMARGI to HEK and HFF cells to yield 361.2 and 355.2 million 2×100 bp paired-end sequencing read pairs, respectively. These resulted in 36.3 (HEK) and 17.8 (HFF) million valid RNA-DNA interaction read pairs, in which approximately 35%, 10%, and 55% were proximal, distal, and inter-chromosomal interactions, respectively (Fig. 1C and *SI Appendix*, Fig. S1A). The proximal and distal interactions were defined as the intra-chromosomal interactions where the RNA-end and DNA-end were mapped to within and beyond 200 kb, respectively. Following Sridhar et al. (10), we removed proximal read pairs from further analysis because proximal interactions likely represent interactions between nascent transcripts and their neighboring genomic sequences. Hereafter, we refer the union of distal and inter-chromosomal interactions as *remote interactions*. The rest of this paper only deal with remote interactions.

**Fig. 1.**
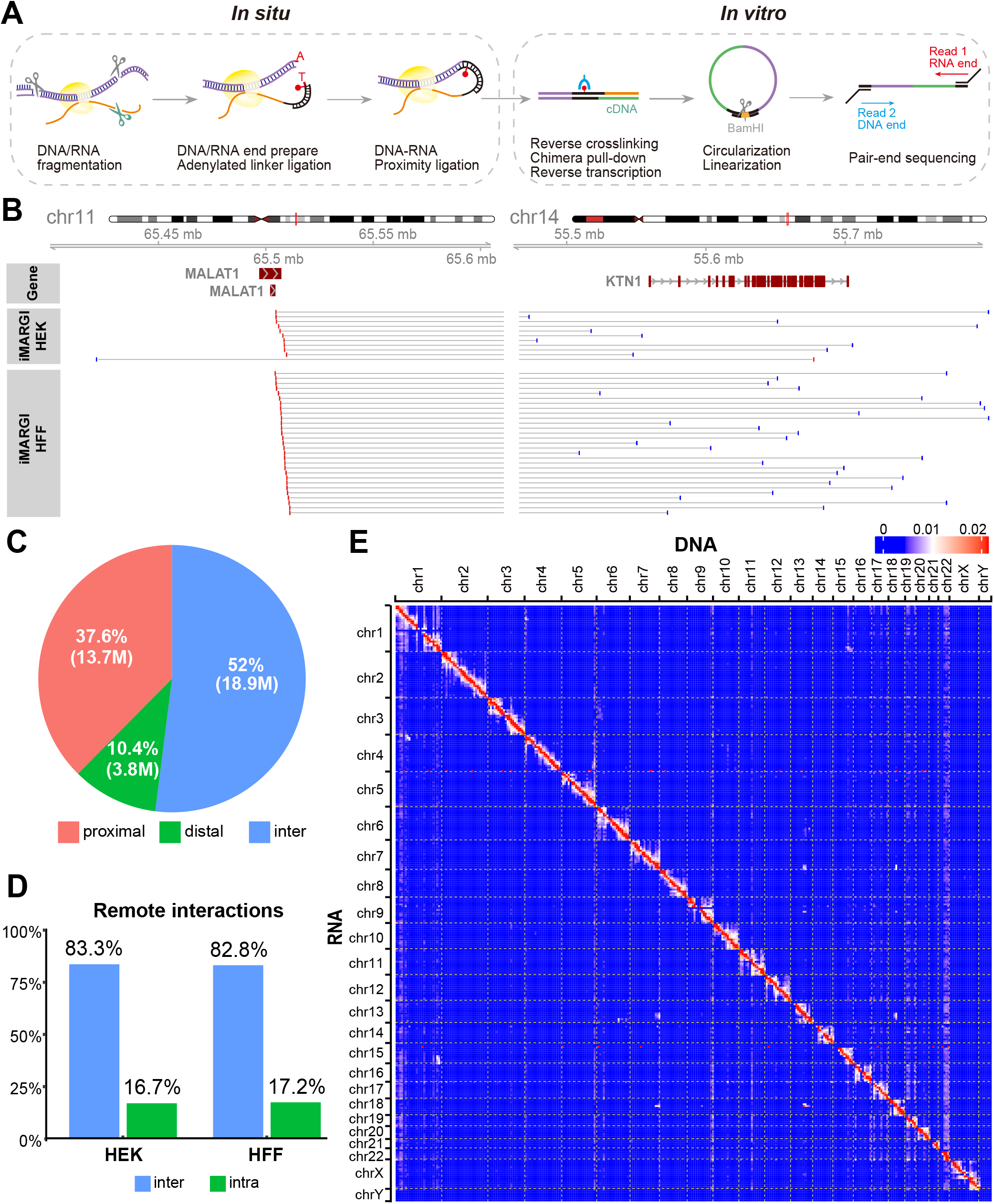
Overview of iMARGI method and data. (A) Schematic view of iMARGI experimental procedure. (B) An example of inter-chromosomal read pairs (horizonal lines), where the RNA ends (red bars) were mapped to the Malat1 gene on chromosomal 11, and the DNA ends (blue bars) were mapped to chromosome 14 near the Ktn1 gene (blue bars). (C) Proportions of proximal, distal, and inter-chromosomal read pairs in collection of valid RNA-DNA interaction read pairs. M: million read pairs. (D) Ratios of inter- and intra-chromosomal read pairs in HEK and HFF cells after removal of proximal read pairs. (E) Heatmap of an RNA-DNA interaction matrix in HEK cells. The numbers of iMARGI read pairs are plotted with respect to the mapped positions of the RNA-end (row) and the DNA-end (column) from small (blue) to large (red) scale, normalized in each row.

Among the remote RNA-DNA interactions, both cell types exhibited approximately 1:5 ratio of intra- and inter-chromosomal interactions (Fig. 1D). The two-dimensional map of RNA-DNA interactions exhibited more interactions near the diagonal line (Fig. 1E and *SI Appendix*, Fig. S1D). Within each chromosome, the number of iMARGI read pairs negatively correlated with their genomic distances (red and blue dots, Fig. 2A).

**Fig. 2.**
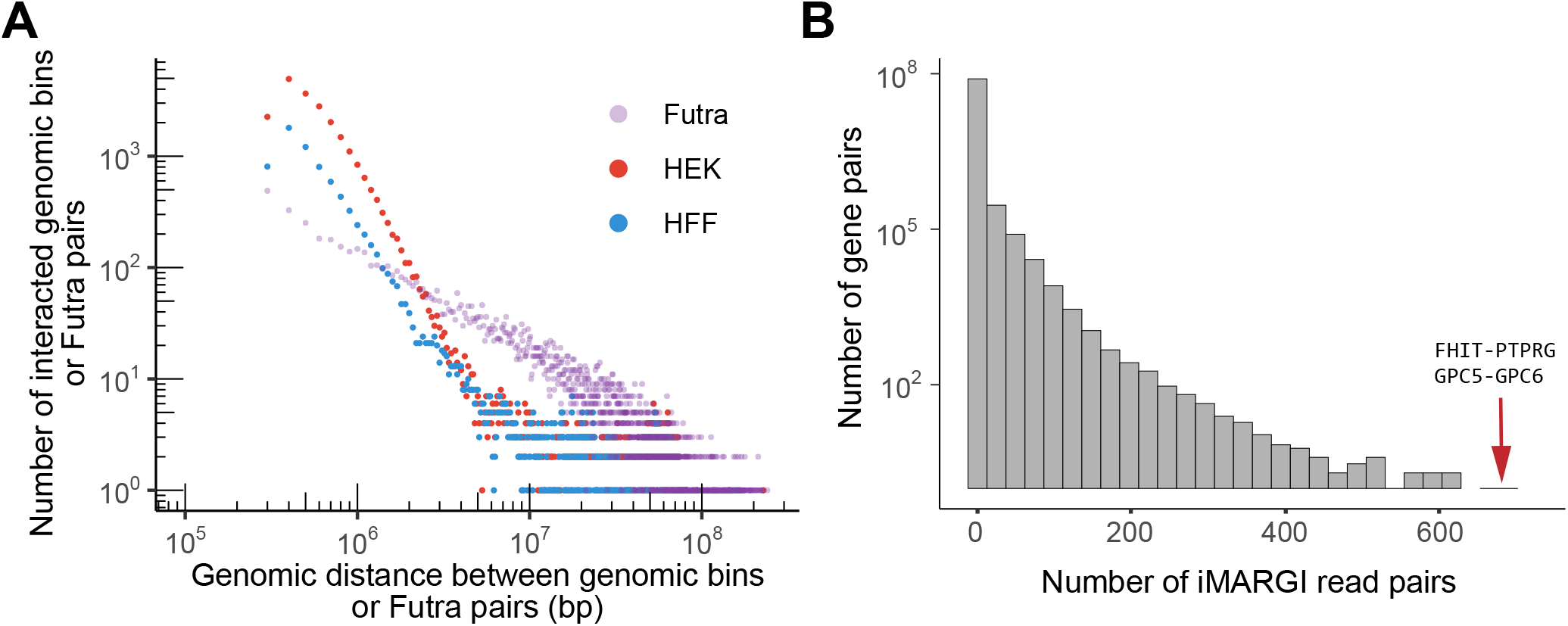
Summary of iMARGI data. (A) The number of genomic bin pairs with 10 or more iMARGI read pairs (y axis) is plotted against the genomic distance between the pair of genomic bins (x axis) in HEK (red dot) and HFF cells (blue dot). For comparison, the number of Futra pairs (y axis) are plotted against the genomic distance (x axis) between the genes involved in the fusion (purple dot). (B) Histogram of the number of iMARGI read pairs of every gene pair (x axis) across all the intra-chromosomal gene pairs with genomic distance > 200 kb. Arrows: the gene pairs with the largest and the second largest numbers of iMARGI read pairs.

### The most significant RNA-DNA interactions co-localized with the gene pairs forming fusion transcripts in cancer

We set off to identify the most significant distal RNA-DNA interactions from the iMARGI data. Excluding extremely abundant non-coding RNAs, such as XIST, the top gene pair with the largest amount of inter-chromosomal and distal iMARGI read pairs in HEK cells was FHIT-PTPRG (Fig. 2B and *SI Appendix*, Fig. S2A). Investigating this gene pair, we found the reporting of FHIT-PTPRG fusion transcripts from kidney, liver, head and neck, lung, and prostate cancers (7). The second largest amount of inter-chromosomal and distal iMARGI read pairs was GPC5-GPC6 (Fig. 2B, *SI Appendix*, Fig. S4 and Fig. S2B). Fusion transcripts from this 2^nd^ ranked gene pair were reported from liver and prostate cancers (7). Notably, 5 of the top 10 gene pairs were reported as fusion transcripts in cancers (1, 7). These findings led us to systematically analyze the relationship between RNA-DNA interactions and fusion transcripts.

### Non-uniform distribution of the RNA pairs contributing to fusion transcripts

We asked whether there is any global characteristic of genome-wide distribution of the RNA pairs that contribute to fusion transcripts. To this end, we subjected the previously reported 16,410 fusion transcripts that were derived from 9,966 samples across 33 cancer types from The Cancer Genome Atlas (TCGA) project to further analysis (4). On average, there were two fusion transcripts per sample (Fig. 3A). The 16,410 fusion transcripts corresponded to 15,144 unique RNA pairs. Hereafter we refer to these gene pairs as “Futra pairs” (<u>fu</u>sion <u>tra</u>nscript contributing RNA <u>pairs</u>). More than 95% (14,482 out of 15,144) of Futra pairs occurred only in 1 sample out of the 9,966 samples analyzed (Fig. 3B). These data confirmed the scarcity of recurrent gene pairs in fusion transcripts (11, 12).

**Fig. 3.**
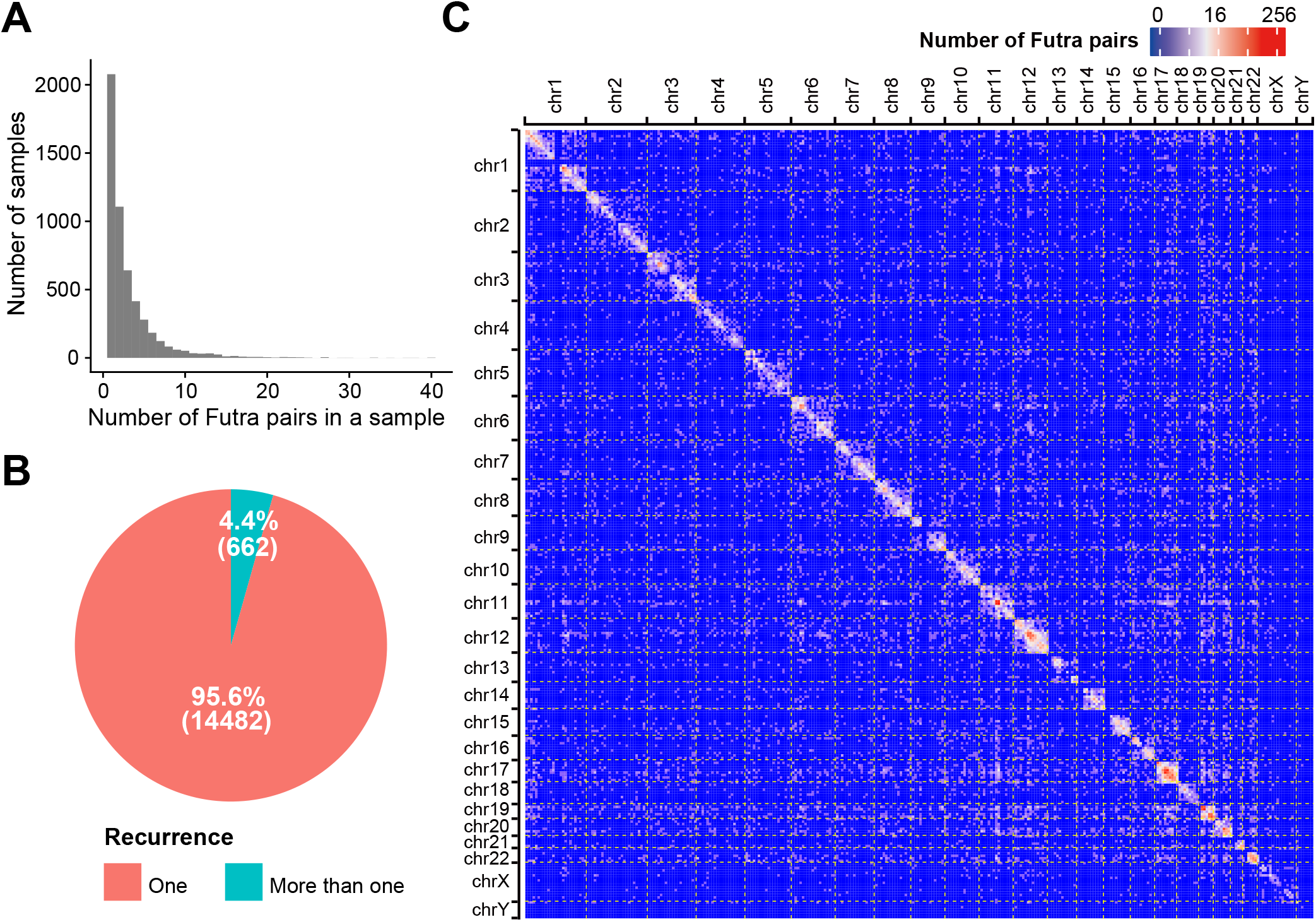
Overview of the 15,144 Futra pairs derived from 33 cancer types. (A) Distribution of the number of detected Futra pairs in each sample across the 9,966 cancer samples. (B) Pie chart of the numbers of Futra pairs that appeared in only one cancer sample (non-recurring Futra pairs, red) and in multiple cancer samples (recurring Futra pairs, green). (C) The genomic distribution of Futra pairs. Genomic coordinates of all chromosomes are ordered on rows and columns. The numbers of Futra pairs of corresponding genomic positions are shown in a blue (small) to red (large) color scheme. Bin size: 10 Mb.

We visualized the frequency of Futra pairs in a two-dimensional heatmap, which we call “fusion map” (Fig. 3C and *SI Appendix*, Fig. S3A). The two-dimensional distribution was not uniform, with more intra-chromosomal than inter-chromosomal gene pairs (odds ratio = 27.91, p-value < 2.2×10^−16^, Chi-squared test). A total of 8,891 and 6,253 Futra pairs were intra- and inter-chromosomal, respectively. Chromosomes 1, 12, and 17 harbored the largest amounts of intra-chromosomal gene pairs *(SI Appendix*, Fig. S3B). Chromosomes 1, 12, 11, 17, 19 contribute to the largest amounts of inter-chromosomal gene pairs *(SI Appendix*, Fig. S3C). Higher density of gene pairs appeared on the diagonal line of the fusion map, suggesting enrichment of gene pairs within chromosomes or large chromosomal domains (Fig. 3C and Fig. 4A). We quantified the relative distances between the Futra pairs. The number of the intra-chromosomal Futra pairs negatively correlated with their chromosomal distance (purple dots, Fig. 2A). Taken together, Futra pairs exhibited non-uniform distribution in the genome, characterized by enrichment of intra-chromosomal pairs and preference to smaller genomic distances.

**Fig. 4.**
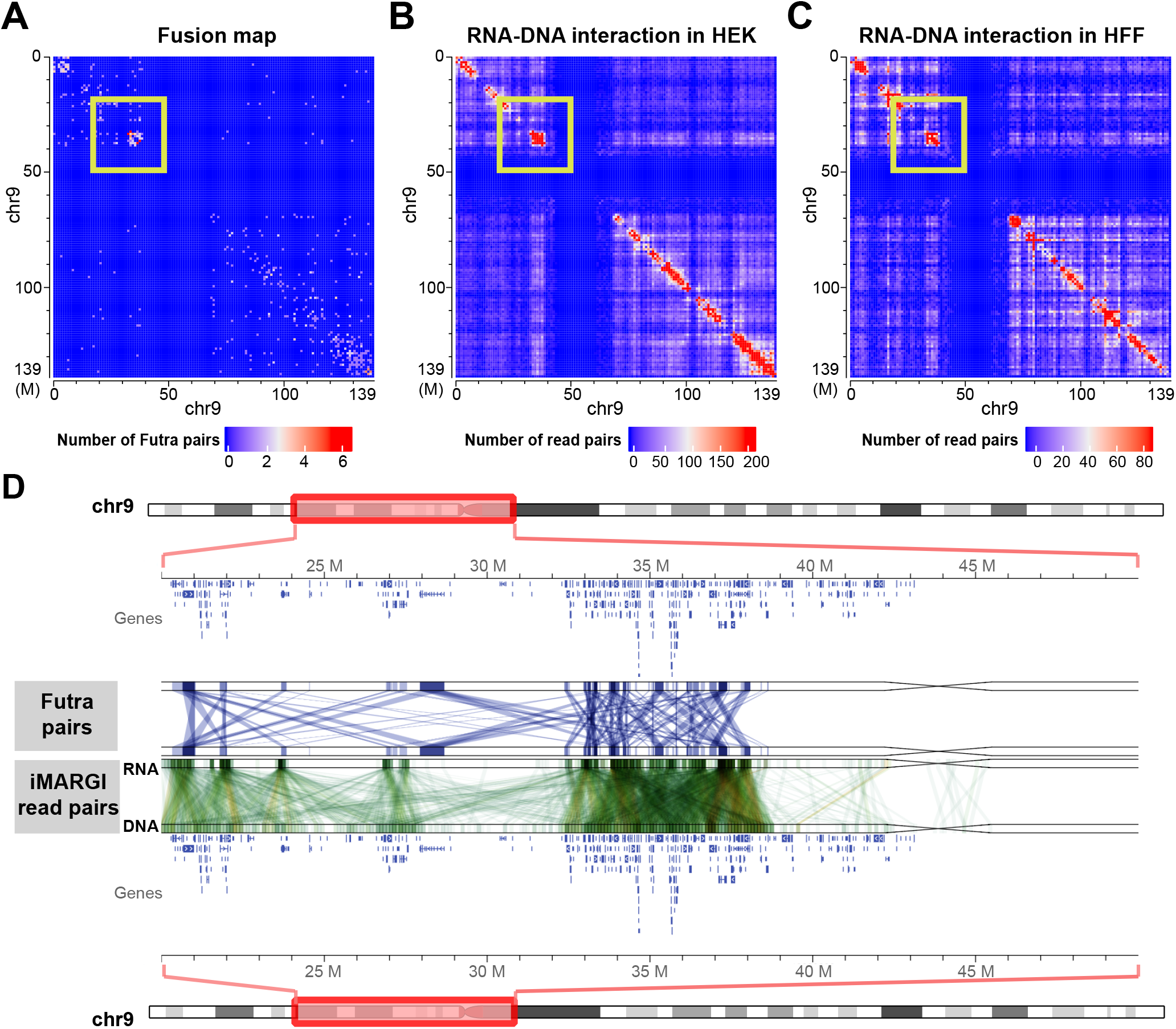
Comparison of Futra pairs and iMARGI read pairs. (A) Distribution of intra-chromosomal Futra pairs on chromosome 9. Genomic coordinates are plotted from top to bottom (rows) and from left to right (columns). (B-C) RNA-DNA interaction matrices in HEK (B) and HFF cells (C). The numbers of iMARGI read pairs are plotted with respect to the mapped positions of the RNA-end (rows) and the DNA-end (columns) on chromosomal 9 from small (blue) to large (red) scale. (D) Detailed view of a 30 Mb region (yellow boxes in A-C). This genomic region and the contained genes are plotted twice in the top and the bottom lanes. In the track of Futra pairs, each blue line linking a gene from the top lane to a gene at the bottom indicates a Futra pair. In the track of iMARGI read pair, each green line indicates an iMARGI read pair with the RNA-end mapped to the genomic position in the top lane and the DNA-end mapped to the bottom lane.

### Differences between the genomic locations of Futra pairs and genome interactions

We asked to what extent Futra pairs may correlate with genome interactions. Forty-one percent (6,253 out of 15,144) of Futra pairs were inter-chromosomal, whereas less than 15% chromosomal conformation capture-derived read pairs were inter-chromosomal (13, 14). The intra-chromosomal Futra pairs exhibited enrichment in large chromosomal domains (Fig. 3C, Fig. 4A and *SI Appendix*, Fig. S4A). These enriched chromosomal domains ranged from approximately 1/10 to 1/3 of a chromosome in lengths, which are approximately 10 - 20 times of the typical sizes of topologically associated domains (TADs) (15). Taken together, Futra pairs exhibited different global distribution characteristics from that of genome interactions.

### Genome-wide co-localization of Futra pairs and RNA-DNA interactions

We asked to what extent Futra pairs may coincide with genome-wide RNA-DNA interactions. We carried out a visualized comparison of the two-dimensional distribution of Futra pairs with that of RNA-DNA interactions (Fig. 3C, Fig. 1E, and *SI Appendix*, Fig. S1D) and observed pronounced similarities (Fig. 4 A-C and *SI Appendix*, Fig. S4 A-C). For example, a set of 34 Futra pairs were enriched in a ~7 Mb region on chromosome 9 (chr9: 32Mb – 39Mb, Fig. 4 A and D). This Futra-pair-enriched region co-localized with a chromosomal region enriched of RNA-DNA interactions (Fig. 4 B-D). Such co-localizations were observed in multi-scale analyses using different resolutions, including 10 Mb *(SI Appendix*, Fig. S4 A-C), 1 Mb (Fig. 4 A-C), and 100 Kb (Fig. 4D) resolutions, as well as at the resolution of individual fusion pairs and read pairs *(SI Appendix*, Fig. S5). Four fusion transcripts, KMT2C-AUTS2, KMT2C-CALN1, KMT2C-CLIP2, and KMT2C-GTF2IRD, were formed between the KMT2C mRNA near the 152-Mb location of chromosome 7 (chr7:152,000,000) and four mRNAs that were approximately 80 Mb away (chr7:66,000,000 – 78,000,000) (Futra pairs, *SI Appendix*, Fig. S5). Correspondingly, a total of 73 RNA-DNA iMARGI read pairs were mapped to KMT2C and the four fusion partners in HEK cells (MARGI, *SI Appendix*, Fig. S5).

We quantified the overlaps between Futra pairs and iMARGI identified RNA-DNA interactions. Among the 6,253 inter-chromosomal Futra pairs, 3,014 (48.2%) overlapped with RNA-DNA interactions in either HEK or HFF cells (odds ratio = 14.1, p-value < 2.2×10^−^, Chi-squared test). Among the 8,891 intra-chromosomal Futra pairs, 7,427 (83.5%) overlapped with RNA-DNA interactions in either HEK or HFF cells (odds ratio = 35.44, p value < 2.2×10^−16^, Chi-squared test). These data pointed to a common feature of cancer-derived Futra pairs, which is their co-localization with RNA-DNA interactions in normal cells.

### Cancer-derived Futra pairs that co-localize with RNA-DNA interaction in normal cells do not form fusion transcripts in normal cells

A model that may explain the co-localization of RNA-DNA interactions and Futra pairs is that RNA-DNA interactions in the normal cells poise for creation of fusion transcripts. Recognizing that this model cannot be tested by perturbation due to the very small likelihood for a fusion transcript to occur in a cancer sample, we carried out two other tests. First, we tested whether the cancer-derived Futra pairs were detectable in normal cells. We re-analyzed the merged RNA-seq datasets of more than 75 million 2×100 bp paired-end read pairs from HEK293T cells (16) and ran STAR-Fusion (17) on these datasets, which reported a total of 8 Futra pairs. None of the previously derived 15,144 Futra pairs from TCGA RNA-seq data were detected in HEK293T cells. In addition, we specifically tested for EML4-ALK fusion transcripts, which were reported in non-small cell lung carcinoma (NSCLC) (18), and there were RNA-DNA interactions between EML4 RNA and ALK genomic locus in HEK and HFF cells (Fig. 6A). Neither PCR nor quantitative PCR analysis detected EML4-ALK fusion transcripts in HEK293T cells *(SI Appendix*, Fig. S6), whereas both assays detected fusion transcripts in a NSCLC cell line (H2228) *(SI Appendix*, Fig. S6). Taken together, these data suggest that, although cancer-derived Futra pairs co-localized with RNA-DNA interactions in normal cells, the fusion transcripts found in cancer are not present in the normal cells.

### RNA-DNA interactions in normal cells are predictive of fusion transcripts in new cancer samples

Next, we tested whether the RNA-DNA interactions in normal cells are predictive of fusion transcript formation in cancer. To this end, we analyzed a validation cohort comprising 96 new lung cancer samples from patients who were not part of the TCGA cohorts. We also analyzed a NSCLC cell line (H2228). RNA was extracted and targeted RNA sequencing (RNA-seq) was carried out with Illumina’s TruSight RNA Pan-Cancer Panel. Of these 96 samples, 27 did not yield sufficient RNA for sequencing, whereas the other 69 samples produced a sequencing library and yielded on average 3.9 million uniquely aligned read pairs per sample (Fig. 5A). The TruSight data analysis package reported a total of 42 fusion transcripts from these 69 samples (Fig. 5 B and C). These 42 fusion transcripts included EML4-ALK and FRS2-NUP107 fusion, which were also reported from the 9,966 TCGA cancer samples, as well as 40 new fusion transcripts that were not previously documented. The small amount of recurring Futra pairs between these additional cancer samples and TCGA samples is expected from the small fraction of recurring Futra pairs across the TCGA samples (Fig. 3B).

**Fig. 5.**
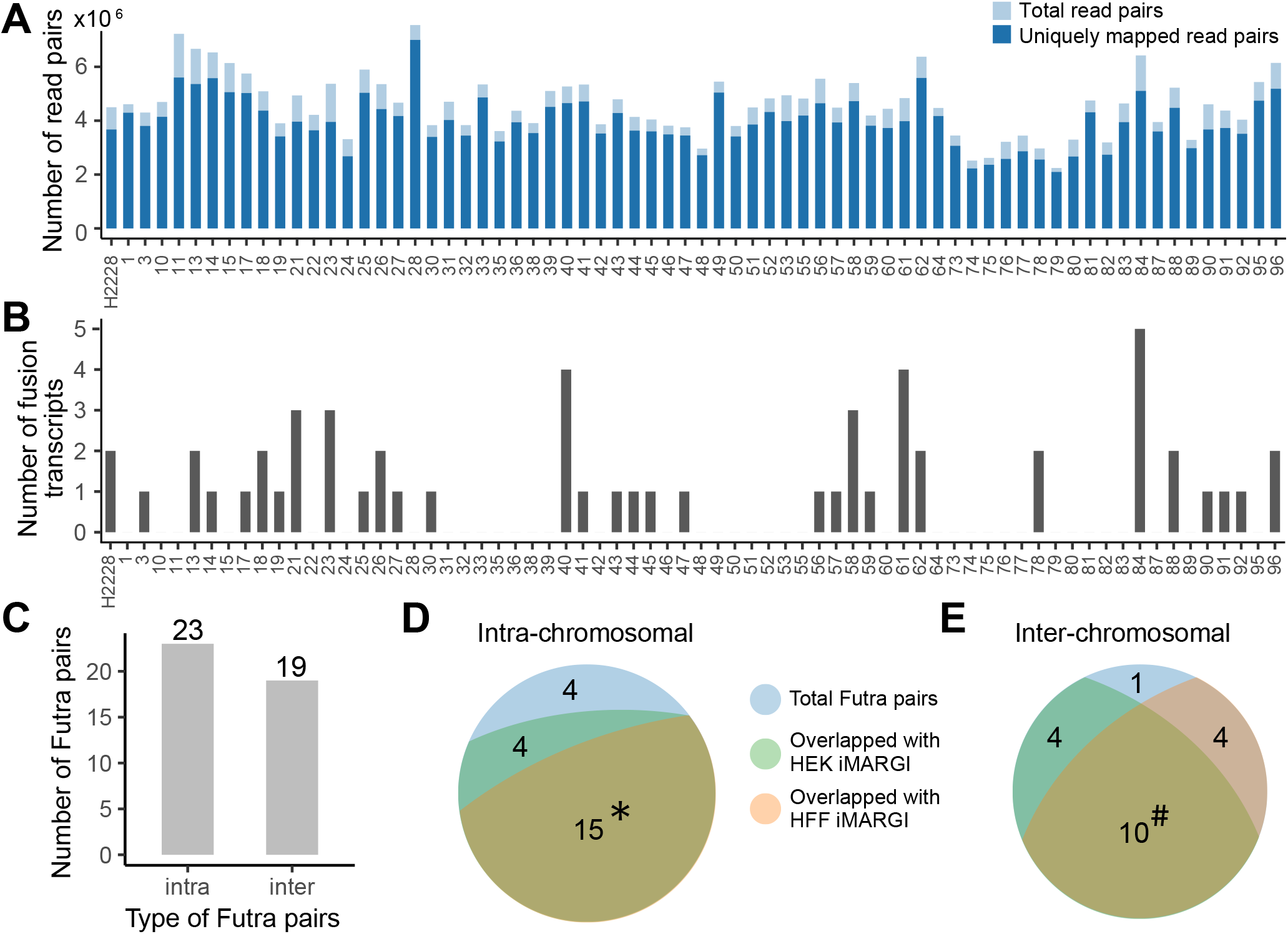
Fusion transcripts detected from the new lung cancer samples. (A) The number of RNA-seq read pairs (light blue bar) and the uniquely mapped read pairs (dark blue bar) of each sample (column). (B) Number of detected fusion transcripts (y axis) in each sample (columns). (C) Numbers of intra- and inter-chromosomal Futra pairs detected from the 69 cancer samples. (D-E) Intersections of these intra-(D) and inter-chromosomal (E) Futra pairs to RNA-DNA interactions in HEK (green) and HFF (pink). *: The 15 intra-chromosomal Futra pairs that overlap with RNA-DNA interactions in both HEK and HFF (yellow-green) are ALK:EML4, RP11-557H15.4:SGK1, FRS2:NUP107, ACTN4:ERCC2, LRCH1:RP11-29G8.3, CUX1:TRRAP, CEP70:GSK3B, NIPBL:WDR70, KCTD1:SS18, CHST11:NTN4, RP1-148H17.1:TOP1, LMO7:LRCH1, NAV3:RP1-34H18.1, LPP:PPM1L, and MTOR-AS1:RERE. #: The 10 inter-chromosomal Futra pairs that overlap with RNA-DNA interactions in both HEK and HFF (yellow-green) are LIN52:PI4KA, LPP:OSBPL6, FCGBP:MT-RNR2, KMT2B:MALAT1, ETV6:TTC3, KTN1:MALAT1, FLNA:MALAT1, COL1A2:MALAT1, COL1A1:MALAT1, and MALAT1:TNFRSF10B.

Among these 42 Futra pairs detected from the validation cohort, 37 (88.1%) co-localized with RNA-DNA interactions in the assayed normal cells, supporting the idea that RNA-DNA interactions in the already assayed normal cells are predictive of Futra pairs in cancer (odds ratio = 106.51, p-value < 2.2×10^−16^, Chi-squared test). We asked if only intra-chromosomal Futra pairs co-localized with RNA-DNA interactions. Nineteen out of the 42 (45%) detected Futra pairs were inter-chromosomal (Fig. 5C), comparable to the proportion (41%) of inter-chromosomal Futra pairs detected from TCGA samples. Eighty-three percent (19 out of 23) intra- and 95% (18 out of 19) inter-chromosomal Futra pairs overlapped with RNA-DNA interactions (Fig. 5 D and E), suggesting that the co-localization of RNA-DNA interactions and Futra pairs were not restricted to intra-chromosomal interactions. Taken together, the co-localization of Futra pairs and RNA-DNA interactions, the lack of cancer-derived fusion transcripts in normal cells, and the predictability to additional Futra pairs in new cancer samples support the model where RNA-DNA interactions in normal cells poise for creation of fusion transcripts in cancers. Hereafter, we refer to this model as the *RNA-poise model*. We call the gene pairs with RNA-DNA interactions in normal cells as *fusion susceptible pairs*.

### RNA-DNA interaction between EML4 and ALK correlates with an RNA fusion without fusion gene in tumor

We tested whether genome re-arrangement is a prerequisite step for the creation of fusion transcripts from fusion susceptible pairs by choosing EML4-ALK fusion transcripts for this test because EML4-ALK is a fusion susceptible pair (Fig. 6A), EML4-ALK fusion transcripts are detected in one of our new tumor samples (Sample #44) (Fig. 6B) and there is an FDA approved diagnosis kit (Vysis ALK Break Apart FISH) based on DNA FISH detection of the EML4-ALK fusion gene. We subjected the remaining tissue from Sample #44 for DNA FISH analysis. None of our 8 attempts yielded any DNA FISH signal in the remaining tissue from either control or ALK probes. We therefore could not ascertain whether there was genome re-arrangement in the only sample with detectable EML4-ALK fusion transcripts.

**Fig. 6.**
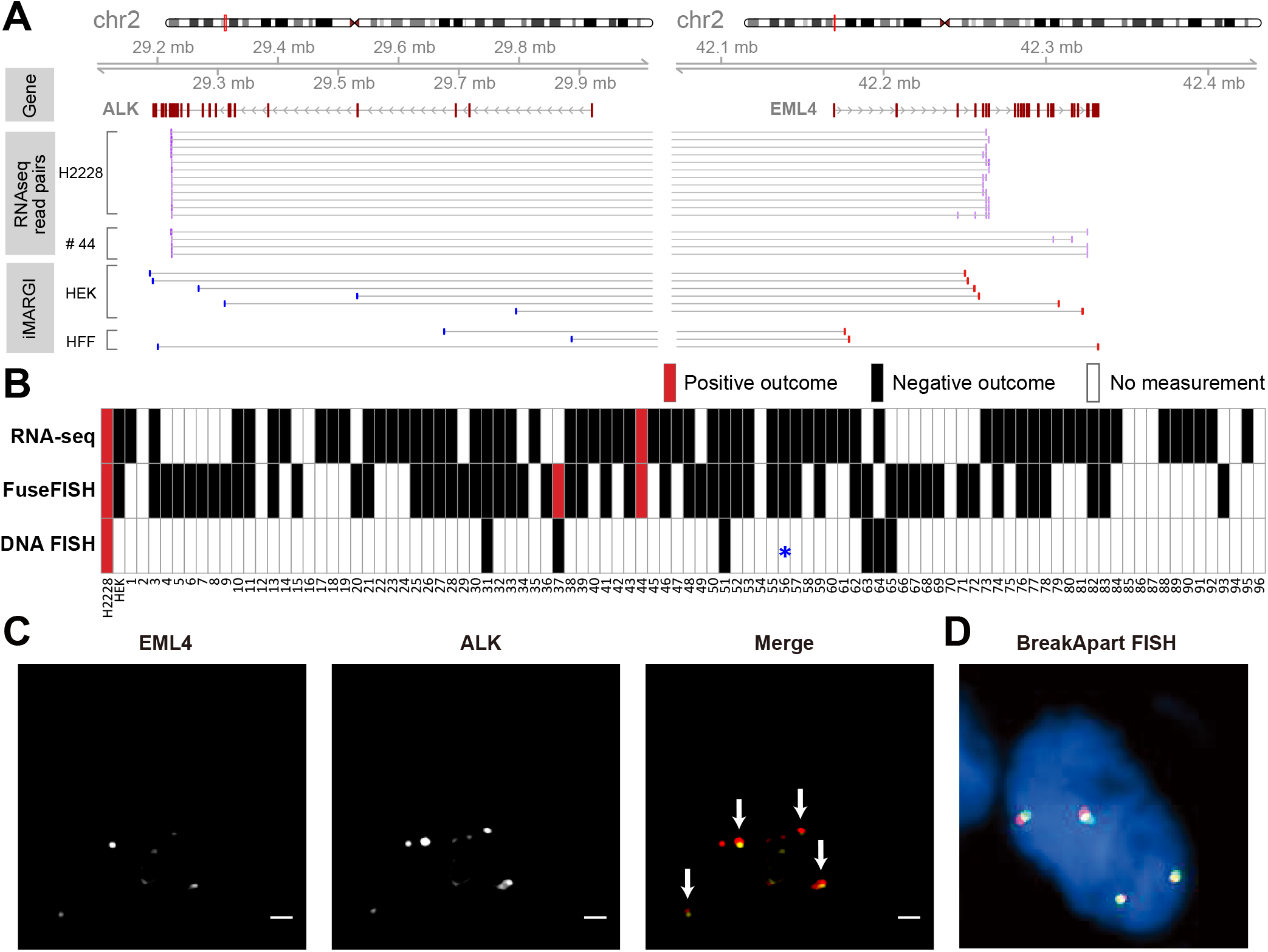
RNA fusion and DNA break. (A) Middle tracks: RNA-seq read pairs (purple bars) aligned to ALK (left) and EML4 (right). Each pair of paired-end reads is linked by a horizontal line. Lower tracks: iMARGI read pairs aligned to the two genes. Red bars: RNA-end. Blue bars: DNA-end. Thin grey lines: pairing information of iMARGI read pairs. (B) Positive (red) and negative (black) outcomes of detection of EML4-ALK fusion transcripts based on RNA-seq and FuseFISH, and DNA FISH (ALK Break Apart FISH, a DNA re-arrangement test) in each cancer sample (column). White box: no measurement. *: Detected a partial deletion of ALK gene without re-arrangement. (C) FuseFISH images of the #37 cancer sample from the EML4 and ALK channels. Arrows: co-localized FISH signals indicating fusion RNA. Scale Bar: 2 *μm*. (D) A representative image of ALK Break Apart FISH, with co-localized red and yellow signals that indicate integral ALK gene without re-arrangement.

In order to identify other cancer samples that express EML4-ALK fusion transcripts, we re-analyzed our collection of 96 lung cancer samples with FuseFISH, a single-molecule fluorescence in situ hybridization (sm-FISH) based method for the detection of fusion transcripts (19, 20). We carried out quantum dot-labelled sm-FISH (21) by labelling EML4 and ALK transcripts with quantum dots at 705 nm and 605 nm, respectively *(SI Appendix*, Fig. S7). The FISH probes were designed to hybridize to the consensus exons shared among all 28 variants of EML4-ALK fusion transcripts that have been identified to date (22). Following prior literature (19, 20), fusion transcripts were detected by the co-localized sm-FISH signals targeting EML4 and ALK transcripts. In a positive control test, an average of 12 co-localized sm-FISH signals per cell were detected in a total of 22 H2228 cells *(SI Appendix*, Fig. S8), that were known to express EML4-ALK fusion transcripts *(SI Appendix*, Fig. S6) (23). In contrast, HEK293T cells exhibited on average zero co-localized signals per cell from 19 cells *(SI Appendix*, Fig. S8), consistent with the lack of such a fusion transcript in HEK293T cells *(SI Appendix*, Fig. S6).

In our collection of 96 tumor samples, only 57 had remaining tissues for FuseFISH analysis. These 57 samples included 39 that yielded RNA-seq data and 18 that did not yield RNA-seq data (Fig. 6B). The FuseFISH analysis detected EML4-ALK fusion transcripts in two samples, including Sample #44 which was also analyzed by RNA-seq and Sample #37 which did not yield RNA-seq data (Fig. 6 B and C). To test whether Sample #37 had ALK-related fusion genes, we subjected it together with other 6 randomly selected samples (#18, #51, #56 #57, #63, #65) for DNA re-combination analysis using Vysis ALK Break Apart FISH (15). None of these seven samples exhibited ALK-related fusion genes. More specifically, one sample (#57) failed to generate DNA FISH signals from four attempts; one sample (#56, negative for EML4-ALK fusion transcripts by RNA-seq and FuseFISH analyses) exhibited a partial deletion of the ALK gene, but no sign of ALK-related fusion genes (star, Fig. 6B). The other five samples, including Sample #37, exhibited integral ALK genes (Fig. 6 B and D). Taken together, the lung cancer Sample #37 expressed EML4-ALK fusion transcripts without having an EML4-ALK fusion gene. These data suggest that genome re-arrangement is not a prerequisite step for the creation of fusion transcripts from fusion susceptible pairs. In other words, the RNA-poise model does not require alterations of the DNA.

## Discussion

### Abundance of genome re-arrangement independent fusion transcripts

Our RNA FISH and DNA FISH analyses revealed a cancer sample that contained a fusion transcript without the corresponding fusion gene. Such an example, although not often seen in literature may not be a rare case (11). The lack of reports are likely attributable to the research attentions paid to the other side of the coin, i.e., the fusion transcripts created by fusion genes (2). Indeed, approximately 36 – 65% of fusion transcripts derived from cancer RNA-seq data were attributable to genome rearrangement (Low Pass bars, Figure S1A of (1)). However, this is likely an overestimate because when low quality whole genome sequencing (WGS) data were removed, only approximately 30 – 45% fusion transcripts had corresponding WGS reads (High Pass bars, Figure S1A of (1)). These published results are consistent with the notion that fusion genes do not account for all observed fusion transcripts and suggest the occurrence of fusion transcripts independent of gene re-arrangement.

### The RNA-poise model allows for splicing errors

Fusion transcripts can be created by two processes. The better recognized process is through transcription of a fusion gene, that was created by genome-rearrangement. The less recognized process is by RNA splicing errors, where two separate transcripts were spliced together (trans-splicing) (24). Trans-splicing does not involve genome-rearrangement. A theoretical gap in the splicing error model is that trans-splicing can only happen to two RNA molecules that are close to each other in the 3-dimensional (3D) space; however, except for neighboring genes (11) the chances for two RNA molecules transcribed from distant chromosomal locations to meet in space are small. Therefore, it remains difficult to perceive a biophysical process in which fusion transcripts are created by splicing errors.

The RNA-poise model fills this theoretical gap. The pre-installation of Gene 1’s transcripts on Gene 2’s genomic sequence positions Gene 2’s nascent transcripts spatially close to Gene 1’s transcripts, allowing for the possibility of trans-splicing. Furthermore, the majority of splicing events are co-transcriptional. The availability of transcripts of Gene 1 during Gene 2’s transcription allows for the opportunity of making co-transcriptional trans-splicing.

### Breaking down the RNA-poise model by RNA-DNA interactions

Remote RNA-DNA interactions could be created by at least two means. First, the caRNA can target specific genomic sequences, which could be mediated by tethering molecules (RNA targeting, Fig. 7). Second, the spatial proximity of the genomic sequences in 3D space could bring the nascent transcripts of one gene to the genomic sequence of another gene (RNA confinement, Fig. 7). Both means of RNA-DNA interactions provide spatial proximity between two RNA molecules and thus allows for splicing errors. In addition, the spatial proximity of two genes in the RNA confinement model could enhance the chances of genome re-arrangement of the spatially close genomic sequences and thus creating fusion genes (25). Thus, the RNA-poise model can be regarded as a union of two sub-models depending on the process of RNA-DNA interaction. One sub-model (Targeting-poise, Fig. 7) could only create fusion transcripts by trans-splicing. The other sub-model (Confinement-poise, Fig. 7) could create fusion transcripts by either trans-splicing or creation of fusion genes.

**Fig. 7.**
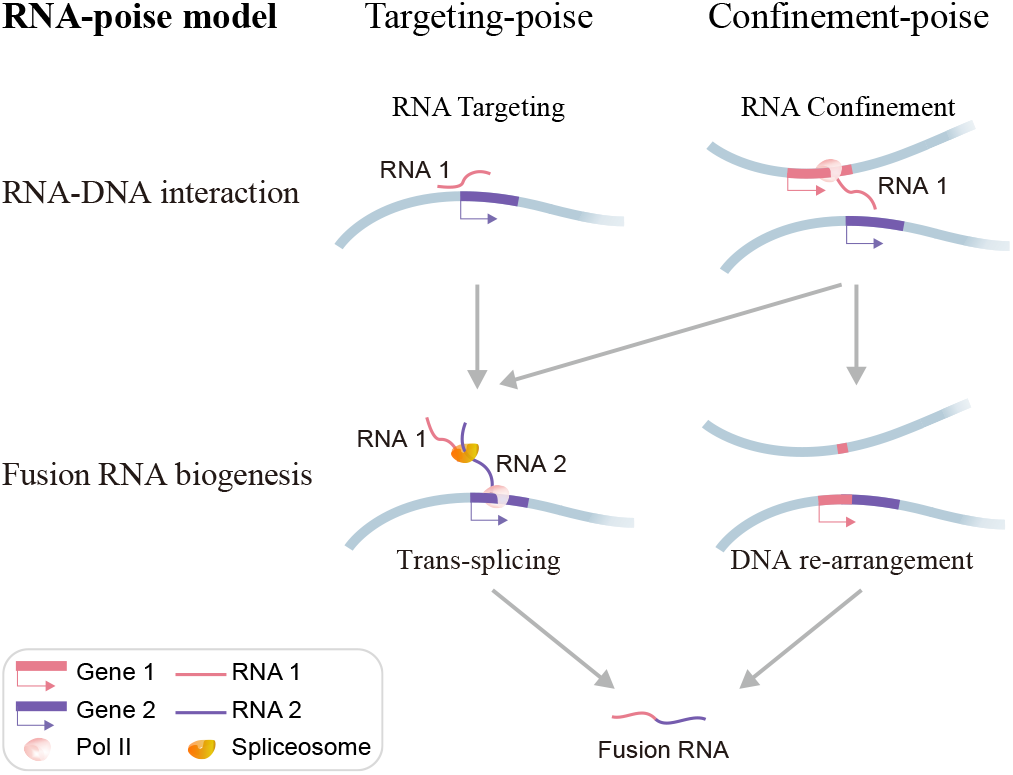
RNA-poise model. In this model, the transcripts of one gene (RNA 1, purple bar) can exhibit spatial proximity to another gene (RNA 2, blue bar) due to tethering (RNA targeting) or spatial proximity of the two genes (RNA confinement). Both cases could enhance splicing errors (grey arrows), whereas the proximity of genomic sequences may also facilitate gene fusion (grey arrow on the right), which subsequently produces fusion RNA.

## Materials and Methods

### Reference genome and gene annotations

Human genome assembly hg38/GRCh38 and Ensembl gene annotation release 84 (GRCh38.84) were used throughout all data analyses.

### Public RNA-seq data

HEK293T RNA-seq datasets were downloaded from NCBI BioProject database (Accession numbers: SRR2992206, SRR2992207, SRR2992208 under the project number PRJNA305831) (16). The three datasets were merged.

### TCGA derived fusion transcripts

The TCGA RNA-seq derived fusion transcripts were downloaded from Tumor Fusion Gene Data Portal (http://www.tumorfusions.org) (26). Tier 1 and tier 2 fusion transcripts were used in our analyses. Genomic coordinates were converted to hg38 by liftOver. Following Davidson et al. (8), Futra pairs within 200 kb on hg38 were removed. The data of Futra pairs were mainly processed using R (27) with Bioconductor packages GenomicRanges (28) and InteractionSet (29).

### Visualization of Futra pairs

Heatmaps of the count matrix were plotted using Bioconductor package ComplexHeatmap (30). Genomic plots of Futra pairs were created with GIVE (31).

## Constructing iMARGI sequencing libraries

### Nuclei preparation and chromatin digestion

Approximately 5×10^6^ cells were used for the construction of an iMARGI sequencing library. Cells were crosslinked in 1% formaldehyde at room temperature (RT) for 10 minutes with rotation. The crosslinking reaction was quenched with glycine at 0.2 M concentration and incubated at RT for 10 minutes. Cells were pelleted, washed using 1×PBS and aliquoted into ~5×10^6^ in each tube. To prepare nuclei, crosslinked cells was incubated in 1 mL of cell lysis buffer (10 mM Tris-HCl pH 7.5, 10 mM NaCl, 0.2% Nonidet P-40, 1× protease inhibitor) on ice for 15 minutes and homogenized with dounce homogenizer pestle A for 20 strokes on ice. Nuclei were pelleted and weighed to estimate the pellet volume (10 mg of nuclei pellet was estimated to be about 10 *μ*L). The nuclei pellet was incubated with SDS buffer (0.5×Cutsmart buffer, 0.5% SDS) at 1:3 volume ratio and 62 °C for 10 minutes with mixing and immediate quenching with a final of 1% Triton X-100. To digest chromatin, the washed nuclei pellet was re-suspended in an AluI chromatin digestion mix (2.3 U/*μ*L AluI (NEB), with 0.3 U/*μ*L RNasinPlus (Promega) and 1× Cutsmart buffer) and incubated at 37 °C overnight with mixing. After chromatin digestion, 1 *μ*L of RNase I (1:10 diluted in 1×PBS) was directly added to the reaction mixture and incubated at 37 °C for 3 minutes to fragment RNA. Nuclei were pelleted and washed twice using PNK wash buffer (20 mM Tris-HCl pH 7.5, 10 mM MgCl2).

### Ligations

To prepare the RNA and DNA ends for linker ligation, nuclei were incubated in 200 *μ*L RNA 3’ end dephosphorylation reaction mix (0.5 U/*μ*L T4 PNK (NEB), 0.4 U/*μ*L RNasinPlus, 1× PNK phosphatase buffer pH 6.5) at 37 °C for 30 minutes with mixing. Nuclei were washed twice with PNK buffer, resuspended in 200 *μ*L DNA A-tailing mix (0.3 U/*μ*L Klenow fragment 3’-5’ exo-(NEB), 0.1 mM dATP, 0.1% Triton X-100, 1× NEB buffer 2) and incubated at 37 °C for 30 minutes with mixing. The same linker sequence as described in the previous MARGI paper was used (10). For *in situ* RNA-linker ligation, nuclei were resuspended in 200 *μ*L ligation mix (38 *μ*L adenylated and annealed linker, 10 U/ *μ*L T4 RNA ligase 2-truncated KQ (NEB), 1× T4 RNA ligase reaction buffer, 20% PEG 8000, 0.1% Triton X-100, 0.4 U/*μ*L RNasinPlus) and incubated at 22 °C for 6 hour and then 16 °C overnight with mixing. After ligation, the nuclei were washed five times with PNK buffer to remove excess free linker. For *in situ* RNA-DNA proximity ligation, nuclei were resuspended in 2 mL of proximity ligation mixture (4 U/*μ*L T4 DNA ligase (NEB), 1× DNA ligase reaction buffer, 0.1% Triton X-100, 1 mg/mL BSA (NEB), 0.5 U/*μ*L RNasinPlus) and incubated at 16 °C overnight.

### Library construction

To reverse crosslinking, nuclei were washed twice with 1× PBS, resuspended in 250 *μ*L of extraction buffer (1 mg/mL Proteinase K (NEB), 50 mM Tris-HCl pH 7.5, 1% SDS, 1 mM EDTA, 100 mM NaCl) and incubated at 65 °C for 3 hours. DNA and RNA were extracted by adding an equal volume of Phenol:Chloroform:Isoamyl Alcohol (pH 7.9, Ambion) followed by ethanol precipitation. The subsequent steps including removal of biotin from non-proximity ligated linkers, pulldown of RNA-DNA chimera, reverse transcription of RNA, DNA denaturation, circularization, oligo annealing and BamHI (NEB) digestion and sequencing library generation were performed as previously described (10). iMARGI libraries were subsequently subject to pair-end 100-cycle sequencing on Illumina HiSeq 4000. The circularization and library construction strategy can phase RNA and DNA ends into Read 1 and Read 2 as shown Fig. 1A, which is the same with MARGI library configuration (10). Since AluI restriction enzyme recognizes “AGCT” sequence and leaves “CT” at the 5’ end of the cut, we expect the first two bases of DNA end (Read 2) to be enriched with “CT”.

## Analysis of iMARGI sequencing data

### Mapping iMARGI read pairs

The detailed iMARGI data processing methods can be found in our GitHub repository https://github.com/Zhong-Lab-UCSD/iMARGI_methods. Briefly, it includes three main steps. First, the read pairs were cleaned by in-house scripts. According to the library construction design, read pairs were filtered out if the 5’-most two bases of their DNA end (Read 2) were not “CT”. Besides, the first two bases of RNA end (Read 1) were removed as they are random nucleotides. Then, the cleaned read pairs were mapped to the human genome (hg38) using bwa mem (version 0.7.17) with parameters “-SP5M” (32). Finally, pairtools (version v0.2.0, https://github.com/mirnylab/pairtools) and in-house scripts were used to parse, deduplicate and filter the mapped read pairs. The valid read pairs that were mapped to genomic locations within 200 kb to each other were defined as proximal interactions, which were excluded from our analysis. GenomicRanges (28) and InteractionSet (29) were used for further analysis of iMARGI data.

### Visualization of iMARGI read pairs

Heatmap of the count matrix were plotted using Bioconductor package ComplexHeatmap (30). Genomic plots of iMARGI read pairs were created with Bioconductor package Gviz (33) and GIVE (31).

### Intersection of iMARGI read pairs and Futra pairs

An iMARGI read pair was regarded as overlapping with a Futra pair when the RNA-end was strand-specifically mapped to the gene body of one gene in the Futra pair and the DNA-end was mapped to the gene body ± 100 kb flanking regions of the other gene in the Futra pair.

## fuseFISH analysis

### Probe design

Oligonucleotide probes were designed to hybridize to exons 2-6 of EML4 RNA and exons 20-23 of ALK RNA. These exons were chosen because they were present in all the observed variations of EML4-ALK fusion genes. These probes were 35-40 nt in lenghs, with similar GC contents and melting temperatures.

### Conjugation of quantum dots to oligonucleotide probes

Oligos were modified on the 5’ end with a primary amino group and a spacer of 30 carbons to minimize steric hindrance of probe-RNA hybridization. These probes were conjugated with quantum dots through the amino group using EDC reaction (34). The probes were subsequently purified with 0.2-μm membrane filtration and 100,000 molecular weight cut-off (MWCO). The retentate of the 100,000 MWCO was subjected with dynabeads MyOne SILANE purficiation to remove any remaining unconjugated probes. A subsequent 0.2-μm membrane filtration was used to remove any final aggregates.

### Hybridization of adherent cell lines

Probe hybridization in H2228 cells was carried out as previously described (19, 21). Briefly, probes were added to the hybridization solution and incubated with the cells at 37 °C overnight. Cells were washed and resuspended in 1× PBS for imaging.

### Hybridization of tissue samples

Probe hybridization in tissue were carried out as previously described (35). Briefly, hybridization solution with probes were added to the surface of parafilm to form droplets. A tissue slice (5-10 *μ*m in thickness) fixed on a glass coverslip was gently placed over the hybridization solution. The mixture was incubated at 37 °C overnight. The tissue was subsequently washed with wash buffer and resuspended in 1x PBS for imaging.

### Imaging and analysis

Cells or tissues were imaged in 1× PBS through wide field fluorescence imaging using an Olympus IX83 inverted microscope at 60 × oil immersion objective (NA=1.4). Image processing was carried out as previously described (36). Briefly, single transcripts were detected using an automated thresholding algorithm that searches for robust thresholds. where counts do not change within a range. Fusion transcripts were determined by searching for co-localization of detection transcripts by overlap between predicted centers within a radius.

### RNA sequencing and analysis

RNA was extracted with Trizol from H2228 cell line and lung cancer tissue samples of the approximate size 3 mm×3 mm×30 *μ*m per sample. RNA-sequencing was carried out using the TruSight RNA Pan-Cancer Panel (Illumina) following the manufacterer’s protocol. All the RNA-seq data, including HEK293T public data, H2228 cell line and lung cancer sample sequencing data, were mapped to the human genome (hg38) using STAR (v2.5.4b) with default parameters (37). Fusion transcripts were called using STAR-Fusion (v0.8.0) (17) requiring both numbers of supporting discordant read pair and junction spanning read larger than zero and the sum of them larger than 2.

## ACKNOWLEDGMENTS

Male hTert-immortalized human foreskin fibroblasts (HFF-hTert-clone 6) are a cell line of the 4D Nucleome Tier 1 cells, provided by Job Dekker lab (https://www.4dnucleome.org/cell-lines.html). This work is funded by NIH R00122368 (Z.C.), HL106579 (S.C.), HL108735 (S.C.), DP1HD087990 (S.Z.), and NIH 4D Nucleome U01CA200147 (S.C. and S.Z.).

**Author contributions**
Conceptualization, Z.Y, N.H., and S.Z.; Methodology, Z.Y, N.H., W.W., and W.C.; Investigation, Z.Y, N.H., W.W., W.C., YJ., J.C., X.H., Z.C., K.Z., S.C., and S.Z.; Writing – Original Draft, Z.Y, N.H. W.W., and S.Z.; Writing – Review & Editing, Z.Y, Z.C., S.C., and S.Z.; Funding Acquisition, S.C. and S.Z.; Resources, K.Z. and S.Z.; Supervision, S.C. and S.Z.
S.Z. is a co-founder of Genemo, Inc., which does not have financial interests with this work. The authors have no financial interests to declare.

